# Logarithmic molecular sampling for next-generation sequencing

**DOI:** 10.1101/420299

**Authors:** Caroline Horn, Julia Salzman

## Abstract

Next-generation sequencing enables measurement of chemical and biological signals at high throughput and falling cost. Conventional sequencing requires increasing sampling depth to improve signal to noise discrimination, a costly procedure that is also impossible when biological material is limiting. We introduce a new general sampling theory, Molecular Entropy encodinG (MEG), which uses biophysical principles to functionally encode molecular abundance before sampling. SeQUential DepletIon and enriCHment (SQUICH) is a specific example of MEG that, in theory and simulation, enables sampling at a logarithmic or better rate to achieve the same precision as attained with conventional sequencing. In proof-of-principle experiments, SQUICH reduces sequencing depth by a factor of 10. MEG is a general solution to a fundamental problem in molecular sampling and enables a new generation of efficient, precise molecular measurement at logarithmic or better sampling depth.

## Introduction

Measurement of molecular abundance is fundamental to basic science and to medicine; it is a classical statistical problem formalized as the problem of estimating the multiplicities *n_i_* of elements *s_i_* in a so-called multiset, and traditionally solved with simple random sampling (SRS). Next-generation sequencing enables SRS of molecules to be performed at high throughput, but in many applications, current or projected throughput is insufficient for addressing critical problems in medicine (e.g. next-generation biomarker detection), chemistry (e.g. high throughput compound screens) and biology (e.g. single-cell sequencing); it is insufficient because SRS suffers from intrinsic limitations when: (i) the cardinality of the multiset is comparable or large compared to the number of measurements taken; (ii) large discrepancies exist between the *n_i_*; (iii) or when precise detection of small changes between the *n_i_* is required. A variety of molecular technologies have attempted to address inefficiencies, including targeted or semi-unbiased enrichment or depletion of a population of molecules (Boone, De Koker, Callewaert, 2018; Hubank & Schatz, 1994). However, these technologies are only semi-quantitative as they compromise quantification of a set of sequences subject to the depletion and require the depleted or enriched sequences to be prespecified.

To overcome these problems, we sought to design an efficient measurement platform with theoretically tractable principles that could be realized in experiment. This led us to develop Molecular Entropy encodinG (MEG), a new general paradigm for molecular sampling that uses computations performed by molecular ensembles to encode the abundance of each species in a sample before measurement, predicts that one can enable experiment-specific sampling designs that only require log or sub-log scale sampling depth compared to SRS while achieving the same measurement precision.

To illustrate the general theory, we start with a simple stylized example (Example 1) of SeQUential DepletIon and enriCHment (SQUICH), a special case of MEG. Consider a tube containing *1* cube and *10^3^* spheres (Fig. 1); these quantities are unknown to the experimenter who wants to estimate them. Suppose there is a physical process that for any number *n* allows up to, but no more than, *n* shapes of each type to be drawn from the tube. SQUICH uses this procedure as follows: in the first round of SQUICH, one object of each shape (cube and sphere) is captured from the tube, tagged with a ``1’’, and placed into a container we call the sampling box. In the second round of SQUICH, up to 1 of each shape is captured and tagged with a ``2’’ and added to the sampling box. Then, up to 9 of each shape are captured and destroyed. In this round, no cubes are captured, as they have all been removed. In the third round, up to one shape is captured, marked with a ‘’3’’, and up to 88 of each shape are destroyed. The numbers of captured and destroyed molecules in the third round satisfy the property that *10^2^* (*1 + 9 + 1 + 89 = 100*) of each shape has been destroyed or captured up to and including this round. In round four, 1 (or 0) of each shape is captured, tagged with a “4”, and up to *898* of each shape are destroyed; in round 5, one more shape is captured and up to 8998 are destroyed, and so on. At the end of 4 rounds, there are 4 cubes and 1 sphere in the sampling box and they are each labeled with the number of the round they were captured in. The order of magnitude of each shape in the original tube can now be estimated with *5* samples without replacement; while SRS requires on the order of *10^3^* samples to make the same inference.

**Figure 1:**
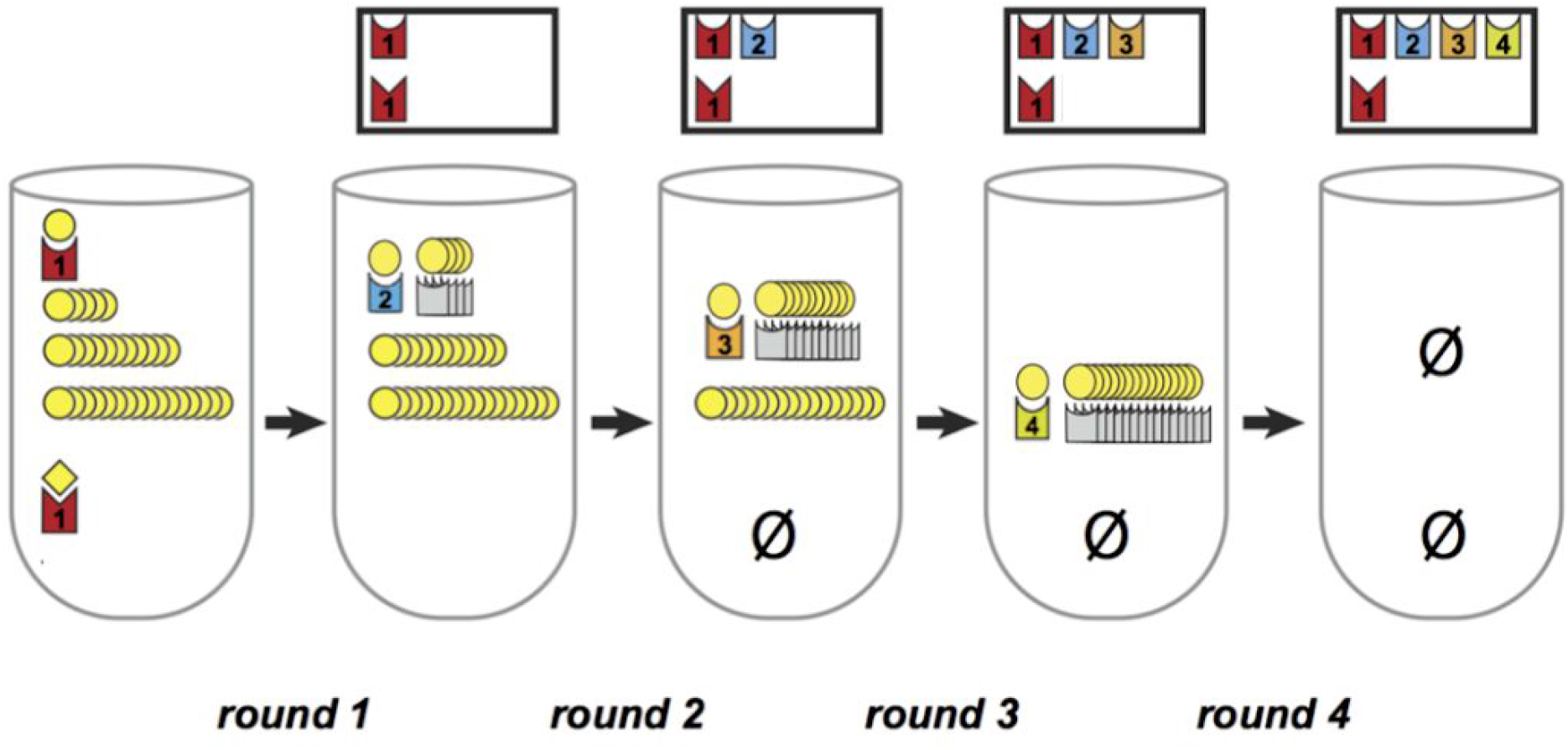
Stylized illustration of SQUICH acting in parallel on spheres and cubes: each interact in parallel with limiting encoders and competitors. When an encoder interacts with a shape, the shape is labeled with that step at which it interacts with the encoder and is brought to a sampling box. The process continues sequentially: encoders are constant amounts while competitor abundance increases (here geometrically). If this geometric increase is in base 10, observation of a tag with the number “4” implies that the original number of molecules in the tube exceeded *10^4^*. The total number of shapes in the sampling box is low, requiring very few samples to fully sample it.

SQUICH is much more general than as presented in Example 1; MEG is yet more general than SQUICH (Methods and Supplement). The numbers in Ex. 1 were arbitrary: the same procedure can operate on, e.g., *10^15^* cubes, allowing a much larger savings in sampling and demonstrating the intuition for why SQUICH enables logarithmic sampling depth compared to SRS.

Informally, three properties enable sampling reductions by SQUICH in Example 1: (1) tagging and removal operating independently on each shape; (2) limiting the number of each shape that is tagged and depleted in each round; (3) sampling only tagged shapes.

The critical properties (1-3) above are fulfilled with nucleic acids replacing the objects of different shapes. Each “shape” in Ex. 1 is replaced by a unique DNA sequence called a target. For each target, sets of DNA oligonucleotides called encoders and competitors that each hybridize with targets are the key to SQUICH. Encoders are libraries of reverse complements of all possible targets which have three critical regions: (1) a region of reverse complementarity to the target; (2) a DNA sequence that is a DNA code representing the round in which the encoder was added to the original tube and; (3) a PCR handle that allows sampling of only targets that extend on encoders. Competitors have the same region of reverse complementarity as encoders, but lack a PCR handle. In each round, targets are hybridized with competitors and encoders, and after hybridization, extend on competitors and encoders (in which case the target is said to be ‘’coded’’) which serve to tag and pull targets into the sampling box as the physical device did for shapes in Example 1 (Figure S4).

As in Example 1, competitors and encoders are added in limiting amounts (*n*) at each step so that removal and/or tagging of no more than *n* of each sequence type can occur in each step. To ensure only coded molecules are sampled, PCR is used as an AND logic gate to selectively sample molecules that are targets AND have extended on encoders (Methods). Competitors can be designed so that targets extending on them can be later retrieved. If targets are in excess of encoders and competitors, the number of targets that extend is limited by the available encoder and competitors. When encoders and competitors are in excess of targets, they compete for binding, which enables estimation of the first significant figures in scientific notation (Results). In addition, the abundance of each competitor and encoder can vary by target as may be desired in certain applications; for example if an experimenter seeks to measure spheres only if they are more abundant than *10^4^* copies, *10^4^* competitors for spheres could be added in the first round.

## Results

SQUICH is simple to embody in experiment and provably enables logarithmic or even sub-logarithmic sampling compared to SRS for precision desired in ubiquitous sequencing applications including estimation of scientific figures formalized in this: **Claim** (Logarithmic sampling with non-filtered round coding): Suppose the abundance of two species are respectively *x_1_10^y1^* and *x_2_10^y2^* with *y_1_ < y_2_* and *x_1_, x_2_* ∈ ℤ^+^ and *0 < p < 1* fixed.There is a SQUICH procedure such that samples *10^y2-y1^* suffice to achieve a probability of detection of at least *p*; a standard result shows SRS requires at least *10^y2-y1^* samples to detect the second species which implies the sampling depth required by SQUICH is logarithmic compared to SRS. See Supplement for proof.

THe proof of the claim shows how SQUICH can achieve more general sampling reductions such as sub-logarithmic rates with super-geometric increases in the number of competitors per round. Simulation tests of SQUICH performance are given in three common application regimes: (1) detection of rare species in the presence of a large background; (2) small fold changes in a complex population; (3) quantification of each species in a population with high dynamic ranges.

To conservatively model SQUICH performance in simulation, we introduce a set of engineered DNA target sequences (which we term "CGA libraries" of length n) consisting of any molecule matching the format [(C/G)A)^n]; CGA libraries are targets behave like the shapes did in Example 1. Competitors and encoders for CGA libraries consist of all reverse compliments of the CGA library with auxiliary sequences that identify them as competitors or encoders (described above, Methods, Tables S4,S5). Equilibrium thermodynamics of CGA are modeled in simulations to include inefficiencies and mismatches in oligonucleotide hybridization when the minimum edit distance between targets is one (Wang & Zhang, 2015; Zhang, Chen, Yin, 2012)(Methods). SQUICH can perform more favorably than in our simulation when targets have minimum edit distance of four or more, a design achieved with sphere packing theory (Conway & Sloane, 2011); that is, CGA codes are a convenient way to explain, model and experimentally embody SQUICH, but SQUICH performance is optimized by different designs of targets. For example, experiments in this paper were performed with oligonucleotides containing degenerate bases (Methods).

Simulation 1 models the “needle in haystack” problem with two species at abundance 10*^x^* where *x*=*15*, with *20* “needle” species at abundance *100*. As predicted by theory, SQUICH robustly identifies all needles across all trials with less than *2000* samples (Fig. 2a). SRS detects at most one needle with 10^11^ samples in 1000 simulations and requires more than 10^15^ samples for the same recall as achieved by SQUICH with 2000 samples (Supplement and basic statistical theory). For *x*= *15*, this implies SQUICH reduces sampling depth by a factor of 10^11^; Table S1, Supplement for other values of *x*.

**Figure 2a:**
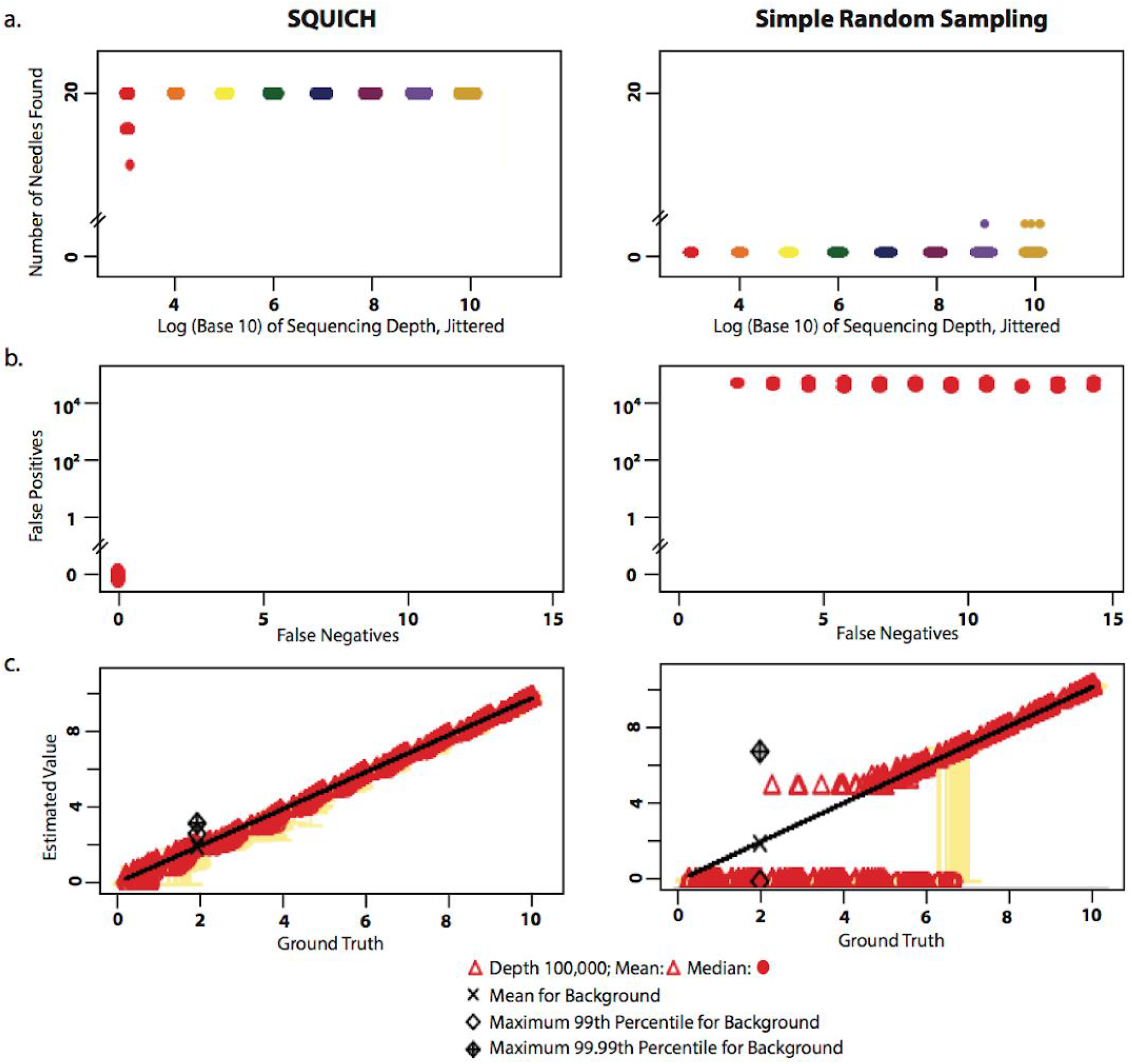
In a background of two species at abundance *10^15^*, SQUICH (L) detects all 20 species with abundance *100* in all simulations with *10^4^* samples; SRS (R) with *10^9^*-fold more reads fails to achieve the same detection yield (1000 trials, x-axis is log*_10_*-scale and jittered). Each sampling depth is depicted in a unique color. **b:** SQUICH enables detection of small fold changes, here 2 fold in 20 species, in a background of *>260,000* species. With *10^5^* samples, all true positives are recovered with 0 false positives. To achieve the same performance with SRS, *10^3^*-fold more samples are required (Supplement) (x- and y-values jittered). **2c:** Detection performance of SQUICH and SRS (100 replicates each) in quantifying species across 10 orders of magnitude. 10 species were assigned an abundance of each value (1:10) *10^0:9^*; and the remaining ~3000 species were set to background level of 100. Yellow bars denote the 25-75th quantiles of measurement for SQUICH and SRS; SQUICH enables detection of small and large molecular abundances across a dynamic range of 10 orders of magnitude with 100,000 samples (top L). At this depth, more than half of species drop-out of SRS sampling and log MSE error is More than 10^10 samples are required by SRS to achieve the same rates of drop-out and log MSE is roughly 2x SQUICH at this depth (Supplement; Table S2).

Simulation 2 tests SQUICH performance where a subset of species (here 20) are 2-fold more abundant than a background of complexity >*260,000*, modeling complexity needed to detect duplication events of > ~10kb in the human genome, or a 2-fold enrichment in a chemical or high throughput pooled CRISPR screen. We designed a statistical estimator for SQUICH to identify species enriched above background (Methods). With this estimator, 10^5^ samples suffice for recovering a median of *18* of the *20* enriched species with 0 false positives (FP) across 1000 trials. To achieve a zero FP rate, SRS requires *10^8^* samples, requiring at least *10^3^*-fold higher sampling depth than SQUICH (Fig 2b). This simulation demonstrates that SQUICH provides flexibility by designing encoder and competitor abundance increases over rounds. Varying competitor and encoder abundance tunes false positive and false negative rates separately, overcoming an intrinsic limitation of SRS where false positives are functionally related to false negatives as a function of sampling depth.

To simulate measurement of native molecules such as RNA or microbial DNA with high dynamic ranges, we modeled each RNA species as a specific CGA code. The molecular biological procedure for converting RNA, DNA or protein to CGA codes is straightforward (Fig. S3; Supplement). In Simulation 3, we test the performance of SQUICH when the distribution of sampled species fills a high dynamic range (*x 10^y^*for *x*=1,..,9 and *y*=0,…,9), as arises in measurement of protein and environmental microbial DNA (Fig. S3). SQUICH fails to detect only 428 of more than 5000 species at a sampling depth by 10^5^ (Fig. 2c); SRS has a drop-out of 3706 species. The log MSE log (Table S3) for SQUICH is lower than SRS at depths up to and including 10^10^. SQUICH performance with 10^6^ samples also exceeds SRS at depth 10^11^; at this depth for SQUICH, a mean of only 4 molecules dropout of sampling.

We also simulated sequencing of a single cells with a dynamic range of transcripts with ~4000 transcripts expressed, and ~1500 transcripts significantly above basal expression which we set to 100 (n=2236), including expression of 10 transcripts at each value *x 10^y^*for *x*=1,..,9 and *y*=0,…,4, and 100 additional transcripts at each level 1:10. With *10^5^* samples, SQUICH has comparable performance to SRS with *10^7^* samples as measured by drop-out (~*2%)* and log MSE (Fig S6; Supplement) a significant improvement over high dropout rates in single-cell sequencing (Vallejos, Risso, Scialdone, Dudoit, Marioni, 2017); SRS at *10^5^* samples has a drop-out rate of roughly 50%, evidence that SQUICH could significantly improve transcript detection in massive throughput single-cell sequencing. In summary, simulation shows that SQUICH exceeds performance of SRS by 100-1000 or more fold in diverse problems including detection of expression of rare species, small fold changes and quantifying species at high dynamic ranges.

SQUICH, as modeled in simulation, can be directly applied to primary biological samples whenever an orthogonal barcode is introduced into the sample, e.g. pooled chemical or genetic screens, with gains in sampling precision illustrated above. To test SQUICH in real next-generation sequencing experiments, we designed a synthetic target library of complexity *2^18^* = 262,144, similar to the CGA code set and manually added a set of individual species ranging from *81_x_* to *80,000_x_* fold over background (Methods, Table S4-5). SQUICH was carried out with 10-fold increases in total molecules in each round, low encoder amounts in the first round and constant encoder amounts in rounds 2-6 (Methods). Six SQUICH libraries prepared with two levels of encoder in round one were sequenced to a mean depth of 2187 reads. Six conventional libraries that model SRS with experimental error introduced during library preparation were sequenced to a mean depth of 19759. In all SQUICH replicates, Pearson and rank correlation between ground truth and estimated abundance exceed all replicates of conventional libraries, despite SQUICH libraries being sequenced at >9 -fold lower depth (Methods, Fig. 3, Tables S4-6).

**Fig 3:**
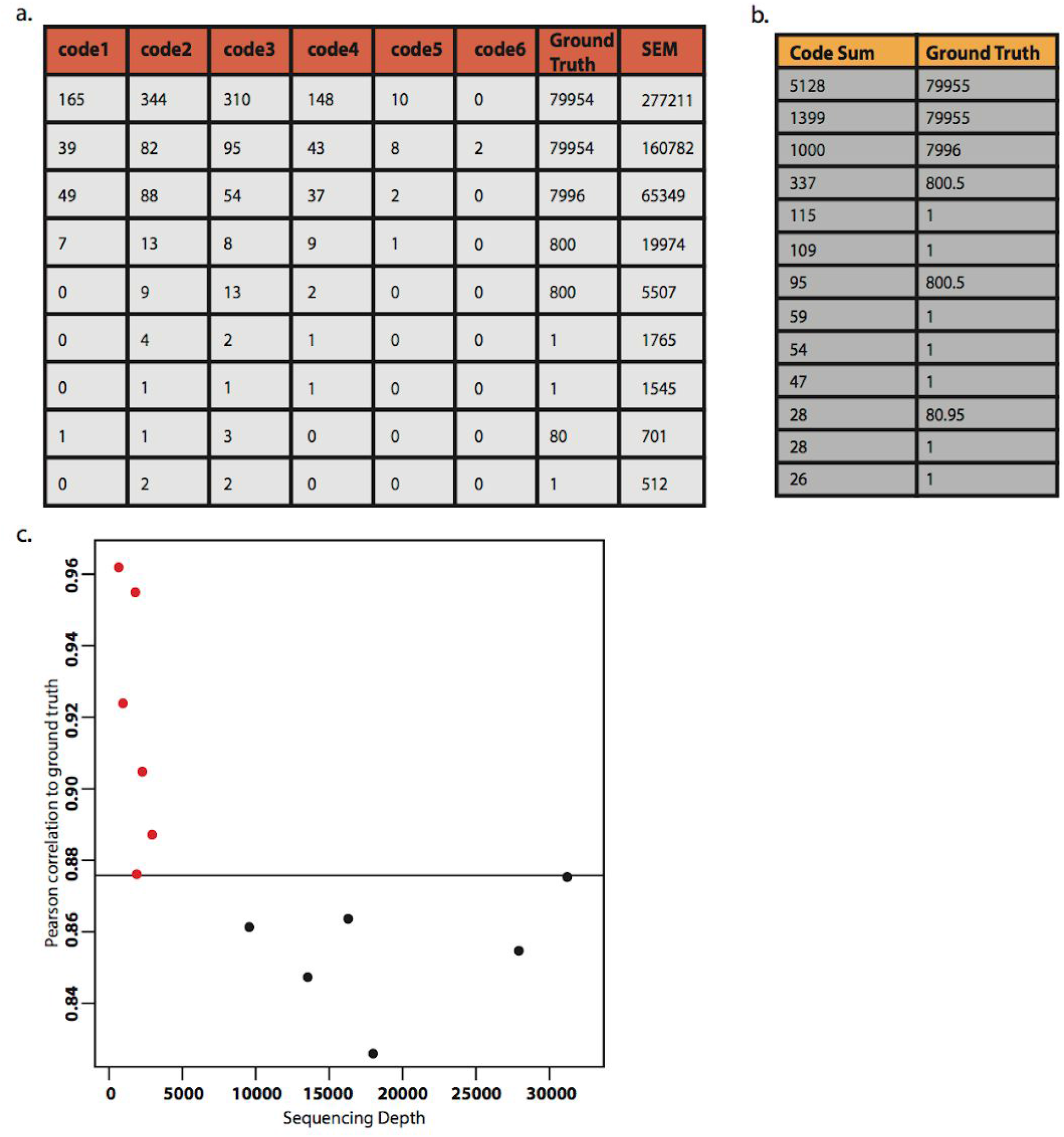
Six SQUICH replicates were sequenced at depths from 1583 to 3305; six conventional sequencing replicates were sequenced to depths of an average of 9 fold greater, from 10345 to 57213 reads. (a) Example of sequencing reads in each code round by SQUICH and SRS (best representative experiment collapsed over two technical replicates for SQUICH and SRS shown); (b) Pearson correlation between estimated counts and ground truth of each SQUICH replicate exceeded each replicate of conventional sequencing although the correlation in one replicate of SQUICH (depth 2407) exceeded the best conventional library (depth 30653) by only ~ .00015.

To control for the high leverage of species with high abundance on correlation values, we used a conservative measure of performance of SQUICH vs. conventional sequencing using a rank based method (Supplement, Methods). 3 of the 6 SQUICH runs sequenced exceeded performance of all 6 conventional sequencing experiments. In addition, 5 out of 6 replicates were statistically significantly more sensitive than SRS with no noise introduced during sequencing (*p*<.05 in 5 out of 6 replicates; *p*=0.138 in one replicate, labeled CH52.03, Table S3). No p-values were significant for conventional experiments. We developed a statistical method to control for variable sampling depths in SQUICH and conventional libraries, and used it to estimate SQUICH efficiency compared to conventional sequencing; it also estimated that proof-of-principle SQUICH experiments achieves a 10x reduction in sequencing depth compared to conventional sequencing.

## Discussion

MEG, and a specific example, SQUICH, is a new framework for quantifying each of a large number (millions or more) species of molecules in a pool, one of the most ubiquitous and important molecular measurement problems today. MEG theory can be applied to any molecular sampling problem, though here we focus on DNA. Small molecules, proteins and RNA can be tagged with DNA sequences, so common assays and screens all reduce to SQUICH, and more generally MEG, measurement. In applications where the sample is limiting, such as biomedical testing, increasing sampling depth is impossible, as sample amplification introduces extra sources of measurement error. In these areas, MEG may be especially important. The flexibility of the sampling distribution provided by MEG expands the scope of statistical algorithms that can be used for estimation. Further, MEG provides key advantages when integrated with modern statistical approaches that use assumptions of sparsity to both improve precision in signal detection and reduce resource cost.

For example, SQUICH could be an ideal platform to measure massive single-cell RNA profiles. To illustrate the design of SQUICH for single-cell RNA-seq, we provide a molecular mapping strategy to combine cell barcodes and gene identity into a single target code as a concise input into SQUICH (Fig. S3). Because this strategy involves hybridization, it has a further unique advantage to improve performance in single-cell applications: multiple target codes can be mapped to the same molecule (e.g. RNA) through hybridization in (Fig S1) with the potential to reduce drop-out, resolve isoforms and overcome 3’ bias or the requirement of a poly-A tail.

We predict that MEG’s design enables even further sampling reductions by providing a platform to convert measurement of nucleic acids into target codes that can be measured by approaches such as compressed sensing, which is not possible achieved with traditional sequencing (Candès, 2006; Cleary, Cong, Lander, Regev, 2017). SQUICH and MEG enable experiment-specific sampling paradigms that lead to future sampling reductions, for example to measure molecules only when their abundance is above a prespecified value. In proof-of-principle SQUICH experiments achieves 10x reduction, we foresee much greater fold reduction by increasing hamming distance between sequences in the pool of targets, competitors and encoders, and increasing purity of oligosynthesis, and by experimental designs that enable specific sampling of only species exceeding or depleted by a prespecified fold. This can be achieved by SQUICH by varying the abundance of each competitor (or encoder) target-by-target, so that for example, either encoders in early rounds are omitted, resulting in only sampling species exceeding a fixed threshold, or increasing encoders in early rounds and decreasing competitors to sample species at low abundance more deeply (unpublished work). In summary, MEG is a new approach for overcoming fundamental limitations in molecular sampling and could enable a new generation of efficient, precise biochemical measurement, from screens to detection of rare species in the blood and single-cell sequencing at an unprecedented resolution, with large numbers of potential variations and platforms.

## Acknowledgements

The authors thank Peter Kim for many useful discussions throughout the project and for an early suggestion of thermodynamic modeling in simulations; Andrew Fire for many useful discussions and critical reading of the manuscript; Peter Wang for useful discussions and generating graphics; Robert Bierman and Julia Olivieri for help with data parsing and critical reading of the manuscript; Eric Freeman for useful discussions, running scripts and generating plots; the Salzman Lab and Rhiju Das for reading of the manuscript. DNA sequencing was performed by the Stanford Functional Genomics Facility, in particular, we thank Xuhuai Ji for running all samples.

## Methods

Caroline Horn and Julia Salzman

### 1 Probabilty models

#### 1.1 Probabilistic formulation of round-coding, definitions

In non-filtered round coding, the *i^th^* SQUICH step is deterministic if *n_i_ > C_i_* + *D_i_* in which case, *C_i_* molecules are coded and *D_i_* are competed out. Because of this, it is useful to introduce the following unique representation of *n_0_* where 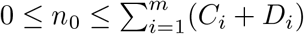which uniquely defines *l*:

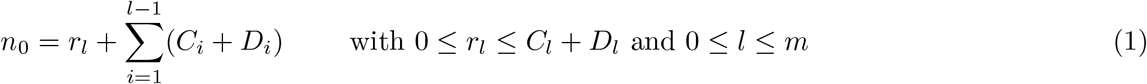

Randomness is only introduced at step *l*, the unique step where encoders and competitors outnumber targets in its round-expansion. Because *C_i_* + *D_i_ > n_i_* in this step, all molecules are coded or competed out.

##### Definition 1.1 (non-filtered round-coding)

*Fix a target sequence and abundance n_0_. For i ≥ 1, and C_i_, D_i_ as the number of encoder i and competitor molecules*.

*Let l be defined by Eqn*. **??**. *Let n_i_ denote the number of free molecules of this target at the end of round i - 1. Let c_i_ and d_i_ be the number of molecules that hybridize and extend on the i^th^ encoder or competitor(respectively). In non-filtered round-coding, for 0 ≤ i ≤ l - 1, c_i_ = C_i_ and d_i_ = D_i_ and*

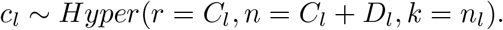

##### Definition 1.2 (Filtered round-coding)

*Fix constants *X*, the discrimination factor and p_t_, p_e_, p_c_, probability of escape of target, encoder and competitor escaping. At step i, if n_i_ molecules of a target are present in round i*,

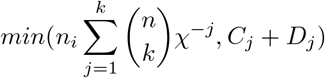

*targets are called ‘conditionally active’. At this round, with probability p_e_ and p_c_ respectively, there are are random number of active encoders and active competitors distributed as B_in_(C_j_, p_e_) and B_in_(D_j_, p_c_) respectively. With probability p_t_, each conditionally active target molecule binds an active competitor and active encoder with equal probability. If bound, a target is considered “active”. Active targets follow the probability distribution in non-filtered round coding*.

### proofs

#### Proof 2.1 (Proof of Claim ??)

*Let*

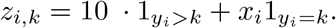

*Then SQUICH transforms the sampling distribution from ratiometric sampling of (x_1_10^y^_1_, x_2_10^y^_2_) to ratiometric sampling of (z_1_,_1_,…, z_1_,y_1_, z_2_,_2_,…, z_2_,y_2_ via the counts from each s_i_-code pair below:*

| sequence | code 1 | code 2 | ... | code $y_2$ |
| --- | --- | --- | --- | --- |
| $s_1$ | $z_{1,1}$ | $z_{1,2}$ | ... | $z_{1,y_2}$ |
| $s_2$ | $z_{2,1}$ | $z_{2,2}$ | ... | $z_{2,y_2}$ |

*Suppose x_1_ = x_2_ = 1, y_1_ =1 and y_2_ is large. Then, to achieve detection of s_1_ with probability p, there exists α_p_ such thatSQUICH requires*

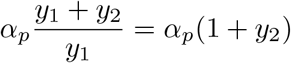

*samples whereas simple random sampling requires* 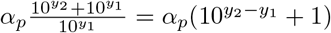.

Geometric progression of 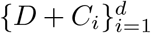is key to encoding scientific digits, as can be seen by a more general choice of *x_i_* and *y_i_* than in the claim. In this case, let *X_i,j_* be the observed counts in the *i, j* cell in the table above.

Further, the expected number of counts in any cell at depth 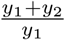 is α. Standard sampling theory implies that

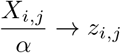

with deviations on the order of 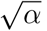. This implies that with high probability, α = 100 suffices for accurate scientific digit estimation.

#### 2.1 Statistical estimators

##### Definition 2.2 (SQUICH EstiMator 1 (SEM1))

*In a SQUICH procedure with d rounds, and satisfying*

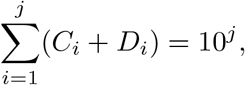

*for 1 <k <m let*

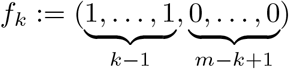

*and e_k_ the k^th^ unit vector*.

*Given data X_1_,…, X_m_ counts observed from each round, let r_k_ be the residual sum of squares from of regressing the vector X on the two orthogonal predictors e_k_ and f_k_, and let*

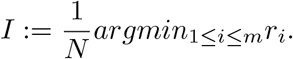

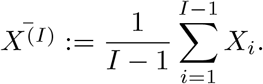

*Then the estimate of nˆ*_0_ *given by SEM1 is*

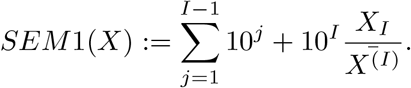

Intuitively, the SEM1 uses a discplined and analytically tractable procedure (regression) to identify the round at which the CGA target is limiting compared to competitors and encoders. The term *X̄*(*I*) serves to normalize observed values of *X_I_* and can be replaced with any estimator of the sequencing depth, or even omitted in applied settings if it suffices to quantify sepcies up to a universal scalar constant. Conditional on correctly determining that round, the SEM1 estimator is then based on the significant digit by estimating a Poisson random variable sampled with a rate proportional to *c_l_*.

In applied settings, we may use the estimator

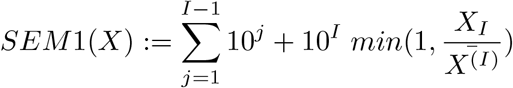

as intuitively, if no coded molecules are observed at round *I* + 1, *SEM* 1(*X*) *≤* 10*^I^*.

It is clear how SEM1 can be extended to arbitrary 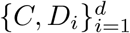 for variable *C_i_*, for example, by defining

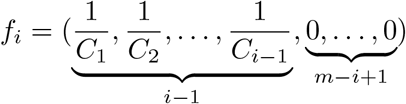

and re-weighting the estimators of *X_I_* and *SEM* 1(*X*) accordingly; for example, the 10^*I*-1^ term in the estimate of *SEM* 1(*X*) in the above estimator derives from the base 10 increase in concentration of the encoder and competitor. In addition, in filtered-round coding, if estimates *C_i_* and *D _i_* of the effective *C_i_* and *D_i_*, each down-sampled by molecular escape, can be computed, they could replace *C_i_* and *D_i_* in the SEM1. The following claim is proved in the Supplement.

##### Claim 2.3 (SEM1 is consistent)

Note that SEM1 could be improved by building strength across samples and using spike-ins such as members of the multiset with ground truth. This allows development of empirical models of the behavior of CGA codes during a SQUICH procedure with increased precision. The full statistical power of spike-ins is a subject for future work. Here, we use spike-ins simply to ensure accurate estimation of tagged molecules in each coding round (see below).

For small sampling depth, SEM1 may have poor performance because it fails to select *I* properly due to high variablility in each coding round. SEM1 could be modified to allow for the values in *f_k_* to take a functional form, such as linear which would result in SEM1 estimating *I* by regression on two predictors. In larger samples, it will be robust to escape of targets as occurs in filtered round coding. Before turning to sampling theory, we introduce a second estimator family (SEM2).

##### Definition 2.4 (SQUICH EstiMator (SEM2))

*In a SQUICH procedure with d rounds, and satisfying*

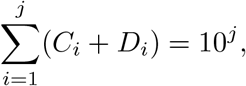

*for 1 <k <m let*

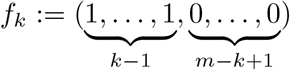

*and e_k_ the ith unit vector*.

*Given data X_1_,…, X_m_ counts observed from each round, the the estimate of nˆ*_0_ *given by SEM1 is*

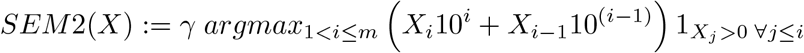

*and γ is a scalar normalizing constant which is a function of sequencing depth*.

As with SEM1, SEM2 has straightforward generalizations to variable encoder and competitor amounts. In addition, in filtered-round coding, if estimates *C*̃*_i_* and *D*̃*_i_* of the effective *C_i_* and *D_i_*, each down-sampled by molecular escape, can be computed, they could replace *C_i_* and *D_i_* in the SEM2. Intuitively, SEM2 will over-estimate *n*_0_ any time a single molecule is sampled at a round that exceeds *log*(*n*_0_). However, SEM2 may have better performance than SEM1 in small sample sizes where SEM1 fails to estimate *I* properly. Other estimators in the SEM2 family replace 1_*X*_*j*_>0 ∀*j≤i*_, a measure that *X*_j_ is not molecular noise but rather represents a true signal that *n_0_ >* 10*^j^*, with an arbitrary function of *X_j_* where *j ≤ i*.

##### Claim 2.5 (Statistical properties of SEM2)

*SEM2 is a consistent estimator of nˆ_0_ and for a geometric progression of 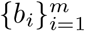 in base b where 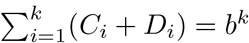, with 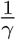 ‖X_*i*_‖∞, in non-filtered round coding*,

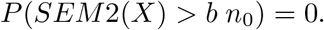

The proof of the claim is straightforward and left to the reader.

Spike-ins are an integral part of each SQUICH experiment. They are designed to bracket the abundance of the range of values to be measured in a SQUICH experiment and will be used to define an estimator that uses empirical behavior of spike-ins to quantify unknown molecular abundance.

##### Definition 2.6 (SQUICH EstiMator (SEM3): interpolation through spikeins)

*Let A be the set of abundances used for spike-ins, S_a_ denote the set of spike-ins at abundance a and n_a_ be the number at abundance a*.

*Let X_i_ be the vector of observed sequencing counts of the i^th^ spike-in. Let X be the vector of observed counts of the target of unknown abundance. Define the following estimator of SEM 3(X):*

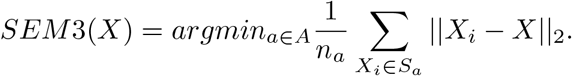

*Then SEM 3(X) is consistent if there exists a ∈ A such that the target is at abundance a*.

It is clear that SEM3 is consistent for values of the parameters that cioncide with with values of the spike-ins and that the bias for *n*_0_ where *n*_0_(*i*) ≤ *n*_0_ ≤ *n*_0_(*i* + 1) is bounded by *|n* _0_(*i*) *- n*_0_(*i* + 1)|.

### 3 Simulation conditions

Parameter regimes were as described in the text and in the parameters in the files multi.squish.data.simulation.r and do.multi.squish.R.

For simulations, SQUICH and simple random sampling were both performed using sampling from the multinomial distribution with fixed sample size as described in the text. Spikeins and species with unknown abundance were both pooled in silico, subjected to SQUICH or left unchanged in the case of simple random sampling trials, and this pool was then sampled.

Simulations that involved SQUICH proceeded as follows. We use the simulation conditions that model the conditions of the “CGA code” as described in the main text. One unit of target, encoder and competitor are each considered to be a mixture of one molecule of each CGA target sequence or its reverse compliment respectively. For each round, the molecules of code used in each round and the cumulative sum of total molecules of competitor and code used up until and including this round were used as parameters to calculate the number of molecules of competitor and code per round (an arbitrary parameterization). In each round, the number of active targets, encoders and competitors were calculated using the following methodology to model noise.

The number of active targets at round *i* was computed as follows:

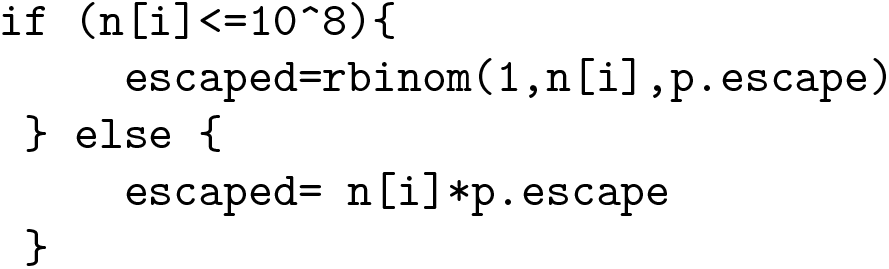

where *p.escape* was a value representing failure of the target to hybridize to either of the compeitor or coder, modeling noise such as binding of oligos to the tube in a molecular biology experiment. Due to numerical conventions in R, if *n*[*i*] exceeded 10^8^, we assigned active targets a value equal to its expectation. The number that “escaped” at this round will always carry through to *n*[*i* + 1], thus a necessary condition that *n*[*i* + 1] = 0 is that *escaped* = 0 above.

We computed active competitors and active encoders molecules in a similar way, but included a term for bias due to off-target binding using a variable thermo.bias. Thermo.bias is an inflation factor modeling imperfect matches between targets and encoder or competitor: with up *k* mismatches with a CGA code of length *n* is modeled as

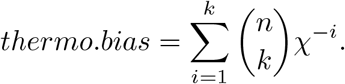

In our simulations, we chose *k* = 3 and *x* as described in the text and below. Given this variable, active competitors and encoders are a random variable with binomial distribution *p* =1 *- p.failure* where *p.failure* is the probability of competitor or encoder escape, and *n* equal to the number of competitor or encoder molecules multiplied by the inflation factor; again, due to numerical conventions in R, we assigned the active molecules to be their expectation if they exceeded 10^8^. A code example is below:

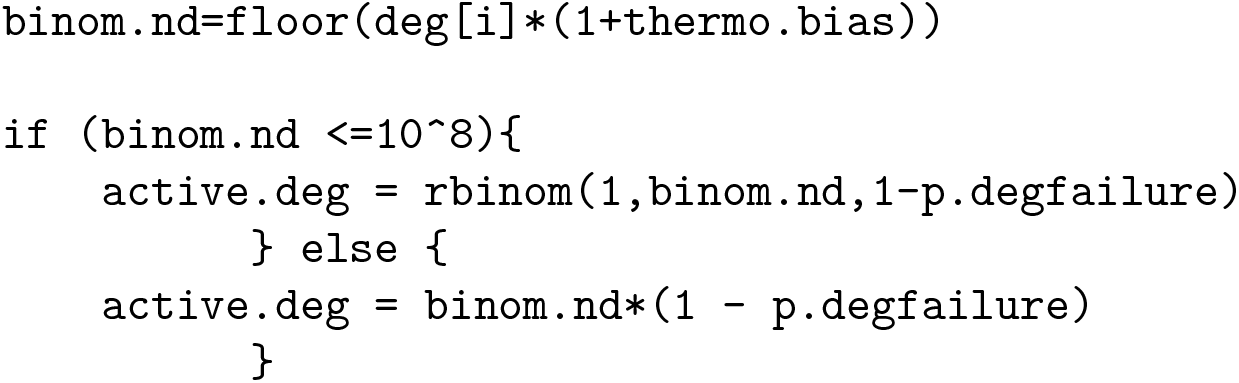

This method treats all molecules as independent, and also inflates the variance (a ‘worse case’) for SQUICH performance by modeling all noise independently.

Given these calculations, if at this round *n*[*i*] exceeds the sum of active encoders or competitors, *n*[*i*] is reduced by the sum of these two numbers, and the number of encoded *n*[*i*] at this round is set to the number of active encoders as calculated above. If *n*[*i*] is less than this number, the number of coded molecules is set to a hypergeometric random variable

> *coded*[*i*] = *rhyper*(*nn* = 1,*m* = *active.code, n* = *active.deg, k* = *n.active*)

and *n*[*i* + 1] is set to the number that escaped.

#### 3.1 Parameters used

##### Experimental Methods

In the experiment without the competitor, a 600 pM solution of the target (disregarding the contribution from the spike ins) was extended using an equimolar pool of the code oligonucleotides (aggregate concentration of 600 pM). Native taq polymerase (Invitrogen) was added in the appropriate buffer and dNTPs, and the solution underwent 5 cycles of denaturing, annealing, and extension. This product was the template for sequencing library 1.

In the SQUICH experiment, the target pool at 600 pM was extended with the first code oligonucleotide (150 pM final) for 5 cycles. After this first extension, a second code (at 150 pM) and a competitor oligonucleotide (at 600 pM) were added and the solution subjected to thermocycling. The addition of code and competitor was performed 5 times with thermocycling between each step. At each step, competitor concentration increased 10-fold to a final concentration of 155 nM. Libraries were prepared for sequencing using standard protocols: sequencing barcodes were added using Phusion polymerase with 15 cycles, and the products were purified via AmpureXP magnetic beads.

The products were then analyzed using the Agilent Bioanalyzer (DNA High Sensitivity) and sequenced on a MiSeq.

Oligo identities are listed in Supplemental Table 1.

Calculations for theoretical performance:

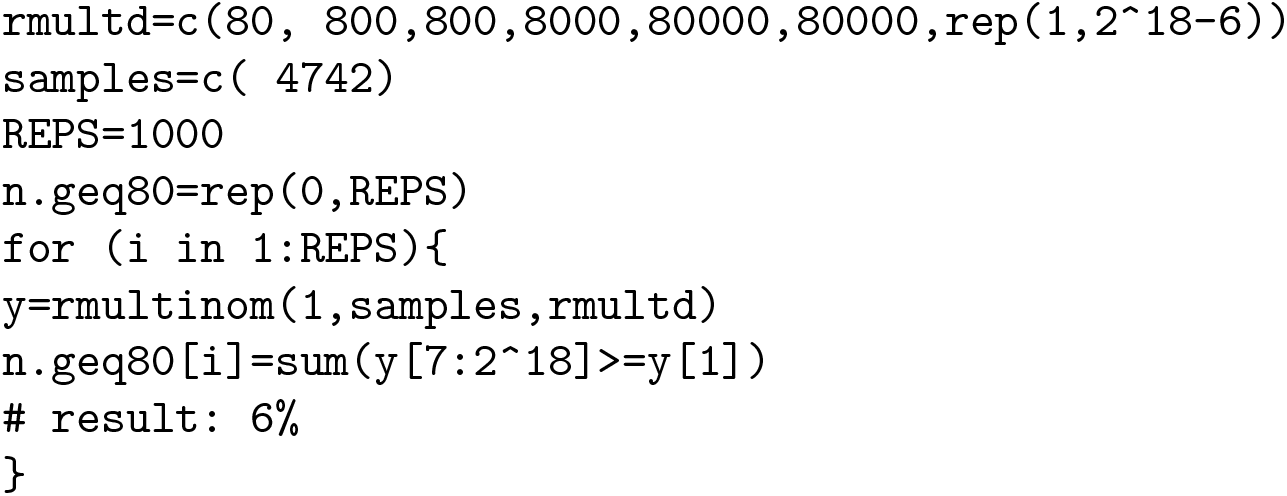

Publication Acknowledgement As with all Stanford Service Centers, credit must be given to the Stanford Functional Genomics Facility for data that results in a publication. If the work done at SFGF produces data resulting in a figure in a publication, you are required to acknowledge the Stanford Functional Genomics Facility in the publication. Further, if SFGF staff members provided significant experimental design, data interpretation, or other intellectual contribution (as evaluated by the PI), then it is expected that these individuals will be coauthors on the publication.

### 4 Supplement

#### 4.1 RNAseq fold change

Here, we systematically profile two performance metrics for SQUICH: (1) detection of low abundance species in a background of high abundance species, ie drop-out; (2) *L*_2_ loss of estimated fold change between estimated abundance and ground truth.

